# Genome-Wide Control of Population Structure and Relatedness in Genetic Association Studies via Linear Mixed Models with Orthogonally Partitioned Structure

**DOI:** 10.1101/409953

**Authors:** Matthew P. Conomos, Alex P. Reiner, Mary Sara McPeek, Timothy A. Thornton

## Abstract

Linear mixed models (LMMs) have become the standard approach for genetic association testing in the presence of sample structure. However, the performance of LMMs has primarily been evaluated in relatively homogeneous populations of European ancestry, despite many of the recent genetic association studies including samples from worldwide populations with diverse ancestries. In this paper, we demonstrate that existing LMM methods can have systematic miscalibration of association test statistics genome-wide in samples with heterogenous ancestry, resulting in both increased type-I error rates and a loss of power. Furthermore, we show that this miscalibration arises due to varying allele frequency differences across the genome among populations. To overcome this problem, we developed LMM-OPS, an LMM approach which orthogonally partitions diverse genetic structure into two components: distant population structure and recent genetic relatedness. In simulation studies with real and simulated genotype data, we demonstrate that LMM-OPS is appropriately calibrated in the presence of ancestry heterogeneity and outperforms existing LMM approaches, including EMMAX, GCTA, and GEMMA. We conduct a GWAS of white blood cell (WBC) count in an admixed sample of 3,551 Hispanic/Latino American women from the Women’s Health Initiative SNP Health Association Resource where LMM-OPS detects genome-wide significant associations with corresponding p-values that are one or more orders of magnitude smaller than those from competing LMM methods. We also identify a genome-wide significant association with regulatory variant rs2814778 in the DARC gene on chromosome 1, which generalizes to Hispanic/Latino Americans a previous association with reduced WBC count identified in African Americans.

## Introduction

The complete genealogy of individuals consists of recent genetic relatedness, such as pedigree relationships of family members, as well as more distant genetic relatedness, such as that due to population structure. In genetic association studies, it is well known that failure to appropriately account for either recent or distant genetic relatedness among sampled individuals can result in spurious association. To address this, linear mixed models (LMMs) have emerged as the standard approach for genetic association testing in samples with population structure, family structure, and/or cryptic relatedness^1-10^. Existing LMM implementations developed for GWAS model the entire genealogy of sampled individuals as a random effect, with the covariance structure of the phenotype specified by a single empirical genetic relationship matrix (GRM)^11-13^. This approach typically provides an acceptable genomic control inflation factor^14^, which is evaluated based on the median of the test statistics across all SNPs genome-wide. However, in the presence of population stratification, previous studies^15,16^ have shown that there may be SNPs for which type-I error rates are not properly controlled, such as those SNPs with unusually large allele frequency differences between populations.

Here, we utilize SNP genotyping data from release 3 of phase III of the International Haplotype Map Project (HapMap)^17^ to demonstrate that existing LMM approaches provide miscalibrated association test statistics when phenotypes are correlated with ancestry. This miscalibration arises due to variation across the genome in allele frequency differences between the populations from which the sampled individuals descend, and we show that it impacts all SNPs genome-wide, not only those with unusually large allele frequency differences. While standard LMM approaches appropriately control type-I error rates at SNPs with typical allele frequency differences, there is systematically inflated or deflated test statistics for SNPs with greater or smaller differences, respectively. Interestingly, we demonstrate that this pattern of test statistic inflation/deflation can occur not only in samples with continental ancestry differences, but also in samples with subtle or fine-scale population structure. Furthermore, the miscalibration of test statistics is observed for LMM methods that estimate variance components once per genome screen, such as EMMAX^3^ and GCTA^13^, as well as those that reestimate variance components for every tested variant, such as GEMMA^8^.

To address the shortfalls of existing LMM methods, we propose a linear mixed model with orthogonally partitioned structure (LMM-OPS) method for genetic association testing of quantitative traits in samples with diverse ancestries. LMM-OPS appropriately accounts for variable population allele frequency differentiation across the genome to provide well-calibrated association test statistics at *all* SNPs genome-wide. With LMM-OPS, genetic sample structure is orthogonally partitioned into two separate components: a component for the sharing of alleles inherited identical by descent (IBD) from recent common ancestors, which represents familial relatedness, and another component for allele sharing due to more distant common ancestry, which represents population structure. LMM-OPS models population structure as a fixed effect by including vectors that are representative of genome-wide ancestry (e.g. principal components (PCs) or admixture proportions calculated from genome-wide data) as covariates, while recent genetic relatedness among individuals is modeled using a random effect, with covariance structure specified by an ancestry-adjusted empirical GRM. An important feature of the GRM used by LMM-OPS is that it is constructed to be orthogonal to the ancestry-representative vectors that are included as fixed effects. This ancestry-adjusted GRM measures the residual genetic covariance among sampled individuals, after adjusting for ancestry, as a way of capturing only recent genetic relatedness. As a result, the ancestry-adjusted GRM and the ancestry-representative vectors represent orthogonal information on sample structure, and LMM-OPS avoids issues of double-fitting information in both the fixed and random effects, which could lead to over-correction of sample structure and a loss of power^6,10^.

We conduct simulation studies to demonstrate that LMM-OPS effectively accounts for complex sample genealogy, including population stratification, ancestry admixture, and familial relatedness, resulting in proper control of type-I error rates at all SNPs, as well as increased power over existing LMM methods for detecting genetic association. We also apply LMM-OPS and the LMM methods implemented in EMMAX^3^, GEMMA^8^, and GCTA^13^ to a GWAS of white blood cell (WBC) count in a sample of 3,551 Hispanic American postmenopausal women from the Women’s Health Initiative SNP Health Association Resource (WHI-SHARe) study^18,19^. The WHI-SHARe Hispanics have complex sample structure, including continental and sub-continental population structure as well as cryptic familial relatedness^20^. Consistent with our simulation study results, the LMM-OPS *p*-values for genome-wide significant SNPs are one or more orders of magnitude smaller than those from the competing LMM methods. Based on our analysis, we replicate^21^ and generalize to Hispanic/Latino Americans a genome-wide significant association with regulatory variant rs2814778 in the Duffy Antigen Receptor for Chemokines (DARC) gene that was previously found to associate with lower WBC count in African Americans^22,23^.

## Materials and Methods

### Standard Empirical GRM

A genetic relationship matrix (GRM), **Ψ**, measures a weighted covariance of genotypes, averaged over all SNPs across the genome, between each pair of individuals. Consider a set 𝒩 of sampled individuals that have been genotyped at a set 𝒮 of SNP genotype markers. A standard empirical estimator^13^ of a GRM that is widely used scales the contribution of each SNP by the sample genotype variance under HWE and has [*i, j*]^*th*^ element

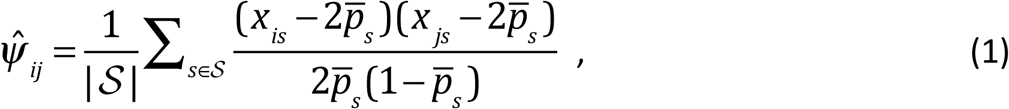

where |𝒮 | is the number of SNPs in the set 𝒮, *x*_*is*_ is the genotype value for individual *i* at SNP *s,* and 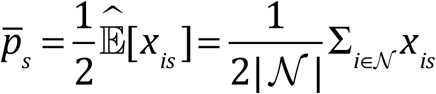 is the sample average allele frequency at SNP *s,* as | 𝒩 | is the number of sampled individuals. The genotype covariance structure captured by 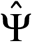, the empirical GRM constructed using the estimator 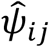 in Equation (1), includes contributions from both distant population structure and recent familial relatedness^24^.

### Construction of an Empirical GRM Orthogonal to Genome-wide Ancestry

Similar to the aforementioned standard GRM, an ancestry-adjusted GRM, **Φ**, also measures a weighted covariance of genotypes, averaged over all SNPs across the genome, between each pair of individuals, however with the covariance is obtained conditional on the genome-wide ancestries of the sampled individuals. Let **V** be an | 𝒩 |×(*k* + 1) matrix whose column vectors include an intercept and ***k*** ancestry-representative vectors (e.g. principal components (PCs) or admixture proportions calculated from genome-wide data). One empirical estimator of an ancestry-adjusted GRM has [*i, j*]^*th*^ element

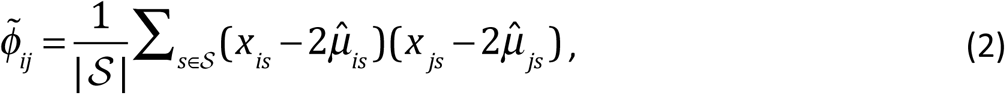

where 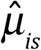 is the *i*^*th*^ element of the vector 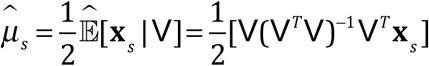 of fitted values from a linear regression of **x**_*s*_, the genotype values for all individuals at SNP *s,* on V. To see that the ancestry-adjusted empirical GRM, 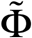 constructed using the estimator 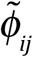 in Equation (2) is orthogonal to genome-wide ancestry, let **R** be an | 𝒩 |×|𝒮 | matrix whose *s*^*th*^column vector is the residual vector from the linear regression of **x**_*s*_ on **V**; i.e. 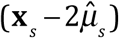. Because the residuals from a linear regression are orthogonal to the predictors, V^*T*^R = **0,** and since the ancestry adjusted empirical GRM can be written as 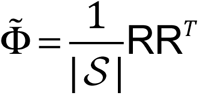, we have that 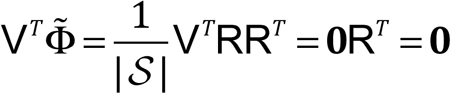, indicating orthogonality of **V** and 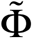. If **V** fully captures the population structure in the sample, then the genotype covariance structure represented by 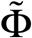 only includes that due to the sharing of alleles IBD from recent common ancestors; i.e. recent familial relatedness^24^.

A potential limitation with 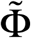, which we refer to as the ‘centered only’ ancestry-adjusted empirical GRM, is that its elements have no meaningful biological interpretation without scaling. To address this, an alternative ancestry-adjusted empirical GRM, 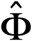, can be obtained using the PC-Relate method^24^, where the [*i, j*]^*th*^ element of this matrix is

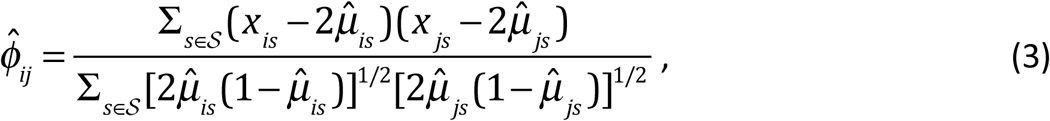

which is an estimator of twice the kinship coefficient for the pair of individuals *i* and *j.* We refer to 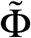 as either the ‘PC-Relate’ or the ‘centered and standardized’ ancestry-adjusted empirical GRM. While 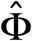 has improved biological interpretability over 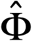, it is no longer strictly orthogonal to the ancestry-representative vectors **V** because the scaling factor in the denominator of Equation (3) depends on *i* and *j.* However, in practice we have found that the scaling factors for each pair of individuals are generally similar, as they are computed as an average across all SNPs in 𝒮, and the elements of 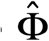 and 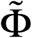 are very highly correlated (see Results), indicating that 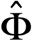 is approximately orthogonal to V.

### Linear Mixed Models for GWAS

A standard linear mixed model (LMM) used in GWAS to test for genetic association at SNP *s*’ ∈ 𝒮 can be written as

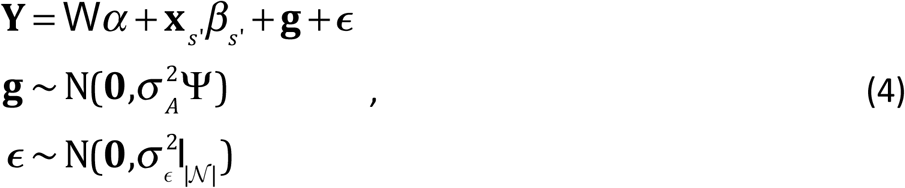

where **Y** is a vector of phenotype values for all individuals, **W** is a matrix of covariates including an intercept, *α* is a corresponding vector of effect sizes, **x**_*s*’_ is the vector of genotype values for all individuals at SNP *s*’, *β*_*s*’_ is the effect size of SNP *s*’, **g** is a random effect that captures the polygenic effect of other SNPs, 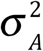 is a parameter that measures the additive genetic variance of the phenotype, **ψ** is the standard genetic relationship matrix (GRM)^25^, **ε** is a random effect that captures independent residual effects, 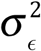 is a parameter that measures residual variance, and I_|𝒩 |_ is an identity matrix. Generalized least squares (GLS) can be used to fit the LMM in Equation (4) and test the null hypothesis that *β*_*s*’_ = 0; however, the overall covariance structure of the phenotype, 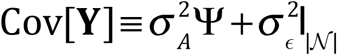, is unknown in practice and must first be estimated. In order to do so, an empirical GRM, such as 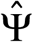 with [*i, j*]^*th*^ element given by Equation (1), is estimated from the available SNP data. Utilizing this empirical GRM in the null model (i.e. the model with *β*_*s*’_ fixed at 0), estimates of the variance components 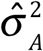 and 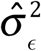 are obtained, typically with restricted maximum likelihood (REML). GLS can then be performed using the estimate of the overall phenotypic covariance structure, 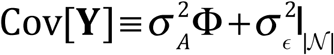.

### The LMM-OPS Model

The LMM-OPS model that we propose has a similar form to the LMM presented in Equation (4), but with the genealogical structure of the sample orthogonally partitioned into fixed and random effects. Population structure is adjusted for as a fixed effect, and recent genetic relatedness is accounted for as a random effect. The LMM-OPS model can be written as

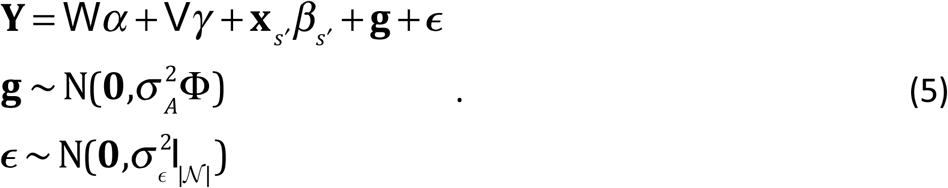

The differences in the LMM-OPS model in Equation (5) from the standard LMM model in Equation (4) are that it includes V, the matrix of ancestry-representative vectors with corresponding effect sizes *γ*, in the mean model to adjust for population structure, and it uses an ancestry-adjusted GRM, **Φ**, that only measures recent familial relatedness, in place of the standard GRM, **Ψ**. Therefore, with population structure modeled as a fixed effect in the mean, the overall covariance structure of the phenotype in LMM-OPS model is given by 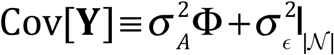. As with the standard LMM in the previous section, GLS can be used to fit the LMM-OPS model and test for genetic association. The procedure is identical, except that 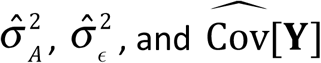 are obtained utilizing an ancestry-adjusted empirical GRM estimated from the available SNP data. Either the ‘centered only’ ancestry-adjusted empirical GRM, 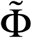 with [*i, j*]^*th*^ element given by Equation (2), or the ‘centered and standardized’ ancestry-adjusted empirical GRM, 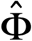, with [*i, j*]^*th*^ element given by Equation (3), can be used for LMM-OPS. Throughout the remainder of this manuscript, unless specified otherwise, we use the centered and standardized ancestry-adjusted empirical GRM when presenting LMM-OPS results.

### Simulation Studies

In all simulation studies, association testing was performed using LMM-OPS, EMMAX, GCTA, GEMMA, and linear regression adjusted for PCs. LMM-OPS included the top PC from PC-AiR^26^ as a fixed effect to adjust for ancestry in the mean model, and it used an ancestry-adjusted empirical GRM constructed with PC-Relate^24^ to account for correlation among genotypes due to recent genetic relatedness. All analyses with EMMAX, GCTA and GEMMA used the default genetic relationship matrices implemented in their respective software to account for sample structure. Details are provided in Appendix A. Throughout the simulation studies, EMMAX and GCTA gave nearly identical results; therefore, only those from EMMAX are presented. Linear regression adjusted for PCs used the top PC from PC-AiR, rather than EIGENSTRAT^27^, to ensure that ancestry was accurately captured and not confounded by pedigree structure^26^.

Simulation studies were used to investigate the impact of variation across the genome in allele frequency differences between populations on association test statistics at null SNPs. Two simulation studies were conducted using samples from two different pairs of HapMap populations: (1) the closely related CEU (Utah residents with Northern and Western European ancestry from the CEPH collection; *n* = 165) and TSI (Toscans in Italy; *n* = 88) populations, which are both European, and (2) the highly divergent CEU and YRI (Yoruba in Ibadan, Nigeria; *n* = 172) populations, which are inter-continental. For each study, we simulated 1,000 replicates of a heritable quantitative phenotype with a mean shift due to an individual’s population membership. To make each replicate of the phenotype 10% heritable, 100 SNPs from chromosome 1 were randomly selected to be causal, each with an effect size chosen based on allele frequency to account for 0.1% of the total phenotypic variability. The effects due to population membership accounted for 18% of the phenotypic variability on average across phenotype replicates in both studies. For each phenotype replicate, the SNPs on chromosomes 2-22, which had no direct causal link to the phenotype, were tested for association. Despite not being causal, SNPs on these chromosomes could be indirectly correlated with the phenotype if they had different allele frequencies in the two populations, resulting in inflated type-I error rates if population stratification was not adequately accounted for.

Additional simulation studies were also carried out to assess the performance of the association testing methods in the presence of ancestry admixture. We simulated genotypes for three separate samples, each with two-way ancestry admixture, but each with a different choice of *F*_*ST*_^28^ (0.01, 0.05, and 0.15) for the underlying populations. An *F*_*ST*_ of 0.01 is a typical value between European populations, such as the CEU and TSI, while *F*_*ST*_ = 0.15 is representative of divergent inter-continental populations, similar to what has previously been estimated between the CEU and YRI populations^29,30^. To generate data sets under each choice of *F*_*ST*_, allele frequencies for the two underlying populations were generated for 200,000 independent SNPs using the Balding-Nichols model^31^, and genotype data at these SNPs were simulated for 3,000 individuals with admixed ancestry derived from the two populations. These SNPs were then split into two disjoint sets of 100,000, which we refer to as Set 1 and Set 2. Each sample included 2,160 unrelated individuals, 120 cousin pairs, and 30 four-generation, twenty-person pedigrees (Figure S1). Individual ancestry proportions for unrelated individuals and pedigree founders were randomly drawn from various beta distributions, and ancestry proportions for pedigree descendants were calculated as the average of their parents’. For each of the three admixed samples, we simulated 1,000 replicates of a quantitative phenotype with 10% heritability and with a mean shift due to an individual’s genome-wide ancestry that accounted for 17% of the phenotypic variability on average. Causal SNPs for generating each phenotype replicate were randomly selected from Set 2. The 100,000 SNPs in Set 1, which had no direct causal link to the phenotype but could be indirectly correlated with it if they were associated with genome-wide ancestry, were tested for association. For each association testing method, sample genealogical structure was inferred using the SNPs in Set 1.

Finally, we performed simulation studies in the admixed setting with *F*_*ST*_ = 0.15 to assess the power of each of the association testing methods to detect causal SNPs. Phenotypes were simulated exactly as for the null SNP studies, but with an additional main effect due to a single causal SNP of interest, *s’*, randomly selected from Set 2. The effect size for this causal SNP was chosen based on allele frequency to account for a pre-specified percentage (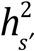= 0.75%, 100%, 1.25%, or 1.50%) of the total phenotypic variability. For each choice of 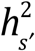, a total of 10,000 phenotype-SNP pair replicates were generated and tested for association. Additional details on how phenotypes and genotypes were generated for all simulations are provided in Appendix B.

### GWAS of WBC Count in WHI Hispanics

The Women’s Health Initiative (WHI) is a long-term national health study focused on identifying risk factors for common diseases in postmenopausal women. A total of 161,838 women aged 50–79 years were recruited from 40 clinical centers in the United States between 1993 and 1998. Detailed cohort characteristics and recruitment methods have been described previously^18,19^. Approximately 17% of participants in this study are under-represented U.S. minority women, and the WHI SNP Health Association Research (WHI-SHARe) minority cohort includes 3,587 self-reported Hispanics who provided consent for DNA analysis. Affymetrix 6.0 genotyping and quality control filtering of these Hispanic-American samples was performed as described previously^32^. Total circulating white blood cell (WBC) count was measured on a fresh blood sample at local clinical laboratories using automated hematology cell counters and standardized quality assurance procedures. Total WBC count was reported in millions of cells per ml, and was log transformed prior to analysis to reduce skewness in the distributions of the phenotypic data. A GWAS of the log-transformed WBC counts measured on women in the WHI-SHARe Hispanic cohort was performed using LMM-OPS, EMMAX, GCTA, and GEMMA. For the LMM-OPS analysis, the first 6 PCs generated with PC-AiR were included as fixed effects to adjust for population stratification, and an ancestry-adjusted empirical GRM calculated conditionally on these PCs with PC-Relate was used to account for recent familial relatedness. The other LMM methods were run with their default settings, filters, and relationship matrices. A total of 616,556 autosomal SNPs were tested for association in the GWAS. Further details are provided in Appendix A.

## Results

### Impact of Variable Allele Frequency Differences on Association Test Statistics

Using the two HapMap based simulation studies, we compared the test statistics obtained from each association testing method for null SNPs. Penalized cubic regression splines were used to find smoothed curves showing the relationship between the absolute value of the allele frequency difference between the pair of populations at SNP *s*, denoted *D*_*s*_, and the mean of the test statistics from each method (Figure 1). Test statistics should follow a 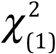 distribution at null SNPs, so the mean of the test statistics for a well-calibrated method should be 1, regardless of allele frequency differences between the two populations. However, in both HapMap studies, the mean of the test statistics from EMMAX and GEMMA increased with increasing values of *D*_*s*_. Test statistics from these methods were substantially inflated (i.e. under-corrected) at SNPs with the largest values of *D*_*s*_ and deflated (i.e. over-corrected) at SNPs with the smallest values of *D*_*s*_. In contrast, the mean of the test statistics from both LMM-OPS and linear regression adjusted for the top PC from PC-AiR showed no relationship with the value of *D*_*s*_. This indicates that including the ancestry-representative PC as a fixed effect in the mean model effectively accounted for the variable allele frequency differences across SNPs. However, since linear regression adjusted for the top PC from PC-AiR did not account for the correlation of phenotypes among relatives, its test statistics were equally inflated across all values of *D*_*s*_ (1.027 on average for the CEU/TSI sample, and 1.032 on average for the CEU/YRI sample). In both studies, LMM-OPS was the only method that provided well-calibrated test statistics for all null SNPs, with the mean of the test statistics near 1 for all values of *D*_*s*_.

**Figure 1.**
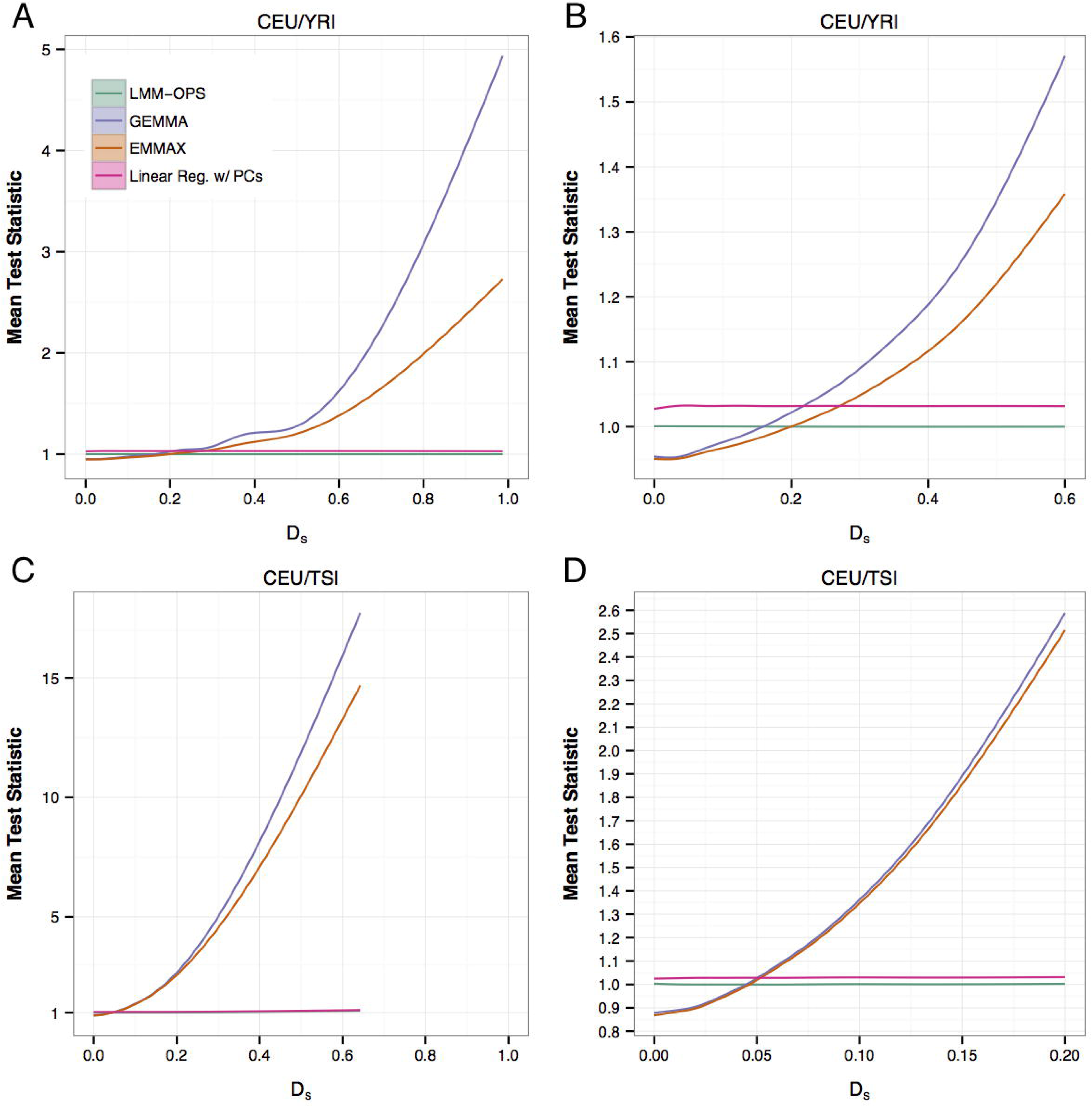
Performance of association methods at null SNPs for a phenotype associated with ancestry in the presence of population structure. Penalized cubic regression splines were used to fit smoothed curves showing the relationship between the absolute value of the allele frequency difference between the two populations at SNP *s, D*_*s*_, and the local mean of the test statistics from each method for the simulations with (A-B) the joint CEU/YRI sample, and (C-D) the joint CEU/TSI sample. The curves shown are the average relationship across all 1,000 simulated replicates (individual points are omitted for visual clarity). The shaded regions show estimated 95% confidence intervals. (A and C) The curves are fit to all SNPs, and the range of *D*_*s*_ is held to [0,1]. (B) The curves are fit to the 98.6% of SNPs with *D*_*s*_ ≤ 0.6. (D) The curves are fit to the 99.9% of SNPs with *D*_*s*_ ≤ 0.2.

Comparing the results from EMMAX and GEMMA for the joint CEU/YRI sample (Figures 1A and 1B) to those for the joint CEU/TSI sample (Figures 1C and 1D), it is apparent that the particular values of *D*_*s*_ for which the mean of the test statistics are either inflated or deflated depends on the pair of populations being analyzed. To further understand this relationship, we investigated the distribution of allele frequency differences across the genome for different pairs of populations by estimating population specific allele frequencies at 1,423,833 autosomal SNPs in the consensus data set for six HapMap populations: the previously mentioned CEU, TSI, and YRI populations, as well as the LWK (Luhya in Webuye, Kenya; *n* = 90), CHB (Han Chinese in Beijing, China; *n* = 137), and JPT (Japanese in Tokyo, Japan; *n* = 86) populations. For each pair of these populations, select quantiles (Table S1) of *D*_*s*_ and its cumulative distribution function across all autosomal SNPs (Figure S2) were calculated. As expected, the distribution of *D*_*s*_ is more concentrated at smaller values for pairs of populations from the same continent (i.e. CEU/TSI, CHB/JPT, and LWK/YRI) as compared to pairs of populations from different continents. Over 90% of SNPs have *D*_*s*_ < 0.1 for the three intra-continental pairs of populations, while at least 49.8% of SNPs have *D*_*s*_ ≥ 0.1 for each of the inter-continental pairs. When SNP *s* has a large (small) *D*_*s*_ value relative to the other SNPs used to construct the GRM, both EMMAX and GEMMA provide an inflated (deflated) test statistic for SNP *s* on average. Therefore, as a consequence of the different distributions of *D*_*s*_, inflation of EMMAX and GEMMA test statistics is observed at smaller absolute values of *D*_*s*_ when jointly analyzing more closely related populations, such as the CEU and TSI, compared to more divergent populations, such as the CEU and YRI.

### Performance at Null SNPs in Admixed Populations

We also compared the test statistics obtained from each association testing method for null SNPs in the simulated admixed populations. Penalized cubic regression splines showing the relationship between the local mean of the test statistics from each method and *D*_*s*_ showed the same patterns as those from the HapMap simulations (Figures 2A-2C). Specifically, LMM-OPS was the only method that provided well-calibrated test statistics for all *D*_*s*_, test statistics from linear regression adjusted for the top PC from PC-AiR were uniformly inflated for all values of *D*_*s*_, and EMMAX and GEMMA provided test statistics that were inflated at SNPs with the largest values of *D*_*s*_ and deflated at SNPs with the smallest values of *D*_*s*_. The distribution of *D*_*s*_ values depended on the *F*_*ST*_ for the pair of populations contributing to the admixed sample (Figure 2D). However, as demonstrated with the HapMap data, the qualitative patterns of test statistic inflation and deflation for each method were the same across all choices of *F*_*ST*_, regardless of the range of *D*_*s*_.

**Figure 2.**
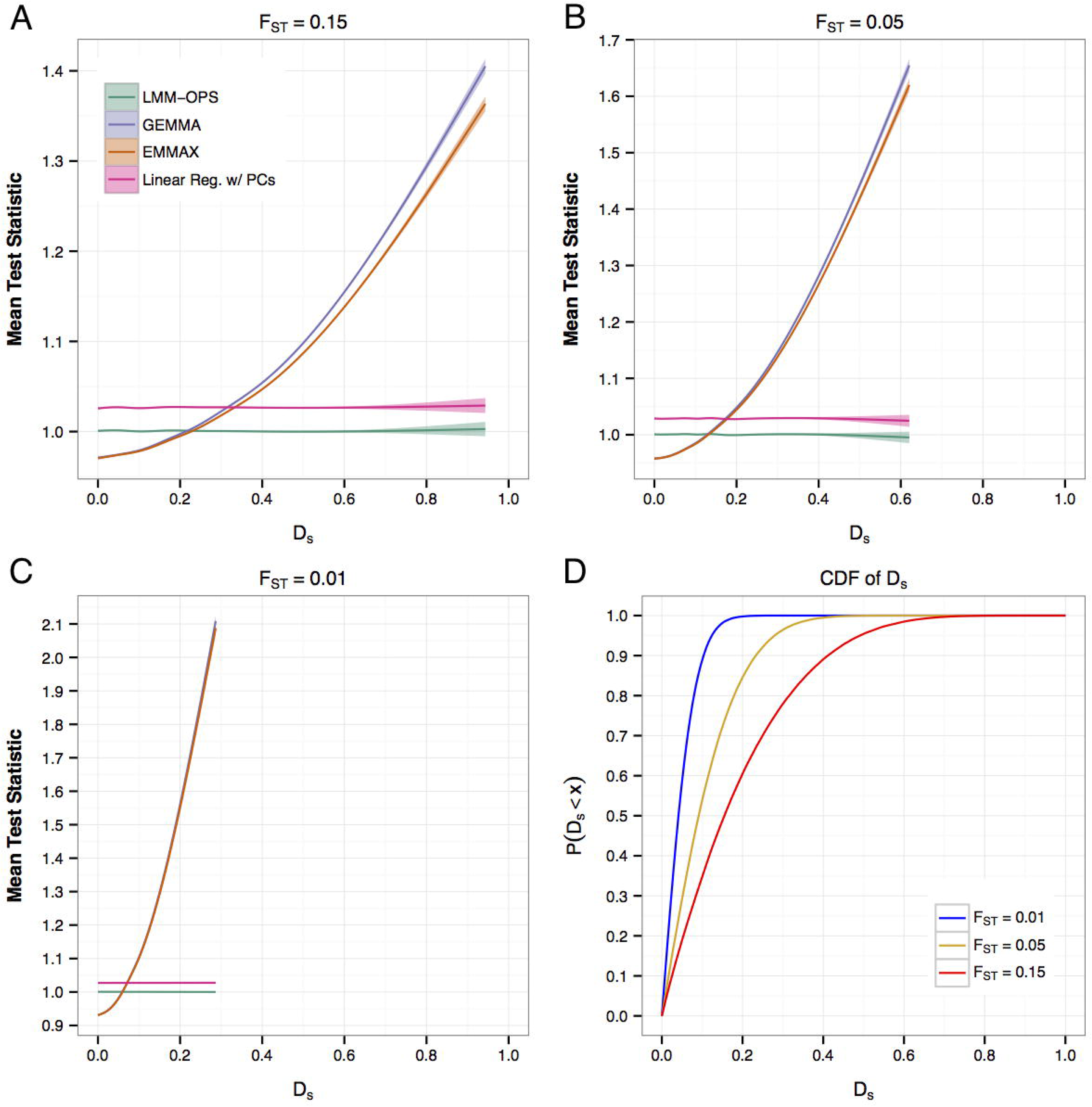
Performance of association methods at null SNPs for a phenotype associated with ancestry in admixed populations. Penalized cubic regression splines were used to fit smoothed curves showing the relationship between the absolute value of the allele frequency difference between the two populations at SNP *s, D*_*s*_, and the local mean of the test statistics from each method for the simulations with admixture from a pair of populations with (A) *F*_*ST*_ = 0.15, (B) *F*_*ST*_ = 0.05, and (C) *F*_*ST*_ = 0.01. The curves shown are the average relationship across all 1,000 simulated replicates (individual points are omitted for visual clarity). The shaded regions show estimated 95% confidence intervals. The range of *D*_*s*_ is kept the same in each panel to emphasize that EMMAX and GEMMA provide inflated (deflated) test statistics at the largest (smallest) values of *D*_*s*_, regardless of its range. (D) Cumulative distribution functions showing the distribution of *D*_*s*_ values for each choice of *F*_*ST*_.

To further examine the performance of each method for null SNPs in the simulation study with *F*_*ST*_ = 0.15, we defined three classes of SNPs based on the magnitude of their allele frequency difference. SNPs were classified as weakly, moderately, or highly differentiated if they were in the first (*D*_*s*_ < 0.07), second/third (0.07 < *D*_*s*_ < 0.28), or fourth (*D*_*s*_ > 0.28) quartile of the distribution of the magnitude of allele frequency differences, respectively. Genomic inflation factors, *λ*_*GC*_, were calculated genome-wide, as well as in each of these three classes of SNPs, for each of the 1,000 simulation replicates (Table 1). The genomic inflation factor is commonly used in genetic association studies to evaluate confounding due to unaccounted for sample structure, where *λ*_*GC*_ ≈ 1 suggests appropriate correction, while *λ*_*GC*_ > 1 indicates an elevated type-I error rate. LMM-OPS was the only method that obtained *λ*_*GC*_ values near 1 genome-wide as well as within all three classes of SNPs. For linear regression adjusted for the top PC from PC-AiR, the average genomic inflation factor was nearly the same (*λ*_*GC*_ ≈ 1.026) genome-wide and within all three classes of SNPs. Interestingly, both EMMAX and GEMMA obtained *λ*_*GC*_ values near 1 when calculated from the median of the test statistics for all SNPs genome-wide, but obtained *λ*_*GC*_ values that were greater than 1 for highly differentiated SNPs and less than 1 for weakly differentiated SNPs.

**Table 1.** Genomic inflation factors, *λ*_*GC*_, at null SNPs for the simulation study with admixture and *F*_*ST*_ = 0.15.

| Method | Genome-Wide | Highly<sup>a</sup><br>Differentiated | Moderately<sup>b</sup><br>Differentiated | Weakly<sup>c</sup><br>Differentiated |
| --- | --- | --- | --- | --- |
| LMM-OPS | 1.000 (0.0002) | 0.999 (0.0004) | 1.001 (0.0003) | 1.001 (0.0005) |
| EMMAX | 1.001 (0.0002) | 1.059 (0.0007) | 0.989 (0.0003) | 0.973 (0.0005) |
| GEMMA | 1.004 (0.0002) | 1.067 (0.0007) | 0.990 (0.0003) | 0.973 (0.0005) |
| Linear Reg. +PCs | 1.026 (0.0006) | 1.026 (0.0007) | 1.027 (0.0007) | 1.027 (0.0007) |

The values presented are the mean (s.e.) across all 1,000 phenotype replicates.
<sup>a</sup> Highly differentiated SNPs: $D_s > 0.28$ between the two populations
<sup>b</sup> Moderately differentiated SNPs: $0.07 < D_s < 0.28$ between the two populations
<sup>c</sup> Weakly differentiated SNPs: $D_s < 0.07$ between the two populations

Additionally, modified QQ plots were generated for each LMM method using the *p*-values for all 100,000,000 null SNPs pooled across the 1,000 phenotype replicates, as well as for the subsets of these SNPs that were highly, moderately, or weakly differentiated (Figure 3). As with the genomic-inflation factors, the QQ plots indicate that LMM-OPS is well calibrated genome-wide as well as in all three classes of SNPs. In contrast, EMMAX and GEMMA appear well calibrated when examining all SNPs genome-wide, but show deviation in the observed *p*-values from those expected under the null when examining each of the three classes of SNPs separately.

**Figure 3.**
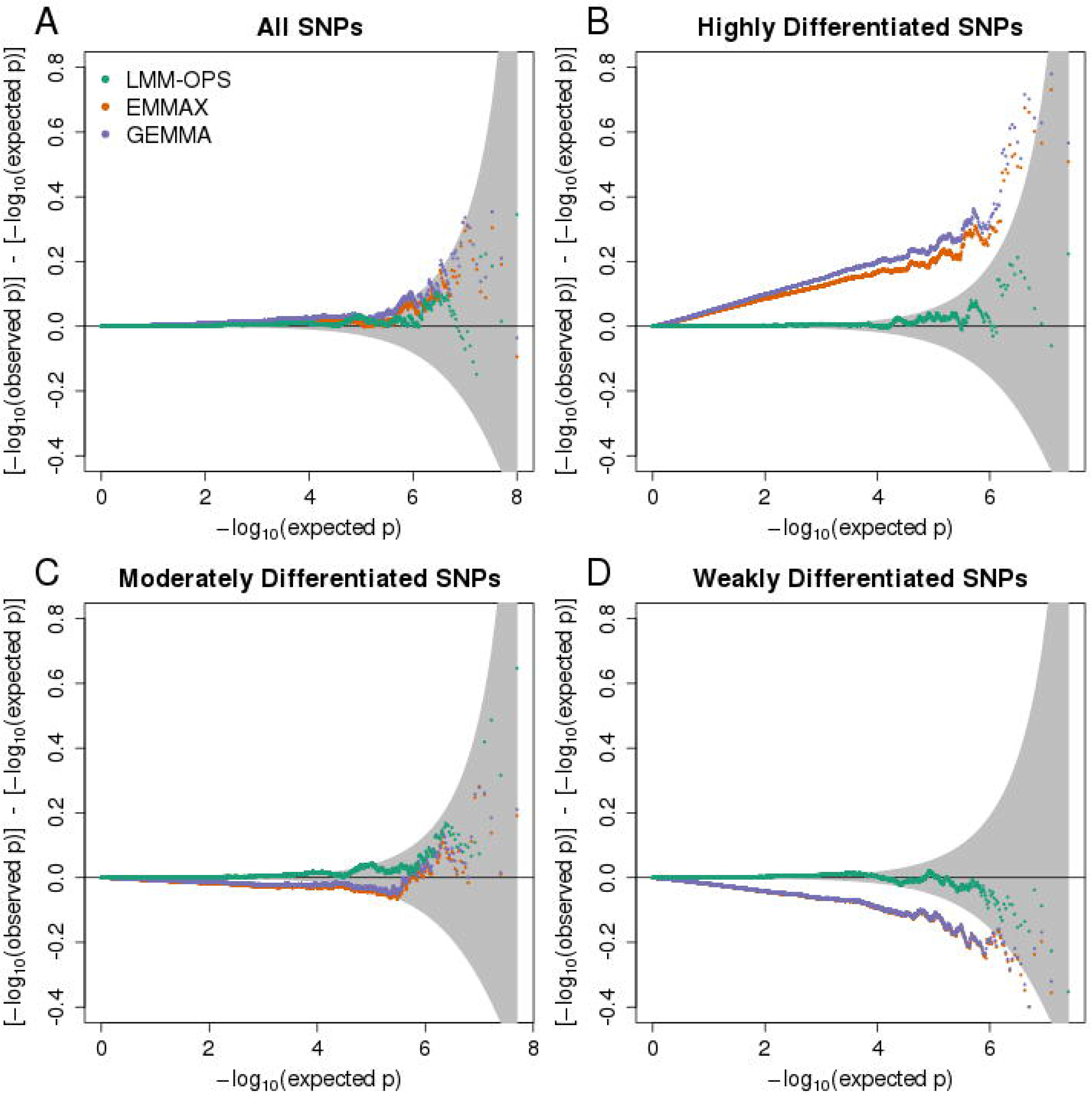
Modified *p*-value QQ-plots for different classes of null SNPs. QQ-plots of *p*-values for (A) all SNPs, (B) highly differentiated SNPs, (C) moderately differentiated SNPs, and (D) weakly differentiated SNPs are presented for LMM-OPS, EMMAX, and GEMMA from the simulation with admixture from a pair of populations with *F*_*ST*_ = 0.15. To more easily see deviation from the null, the y-axis is the difference between the observed and expected –log_10_(*p*-values) rather than the observed. Points above the gray cone indicate inflation, and points below indicate deflation.

### Detection of Causal SNPs

We also performed simulation studies in the setting with *F*_*ST*_ = 0.15 to assess the power of each of the LMM methods. Linear regression with PCs was omitted from these comparisons because it had consistent inflation of type-I error rates across all SNPs. Power to detect causal SNPs with 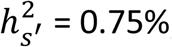, 1.00%, 1.25%, and 1.50% was computed at the genome-wide significance level. *α* = 5 x 10^−8^ across all SNPs, as well as within the highly, moderately, and weakly differentiated classes. When considering all causal SNPs, LMM-OPS had significantly higher power than EMMAX and GEMMA by about 2-3% for each choice of 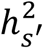 (Figure 4 and Table S2). This difference in power corresponded with LMM-OPS detecting between 2% and 10% more causal SNPs than the other LMM methods. Furthermore, LMM-OPS provided the highest power to detect causal SNPs within each class of allele frequency differentiation. Perhaps surprisingly, this included highly differentiated SNPs, for which EMMAX and GEMMA provide systematically inflated test statistics at null SNPs and have inflated type-I error rates.

**Figure 4.**
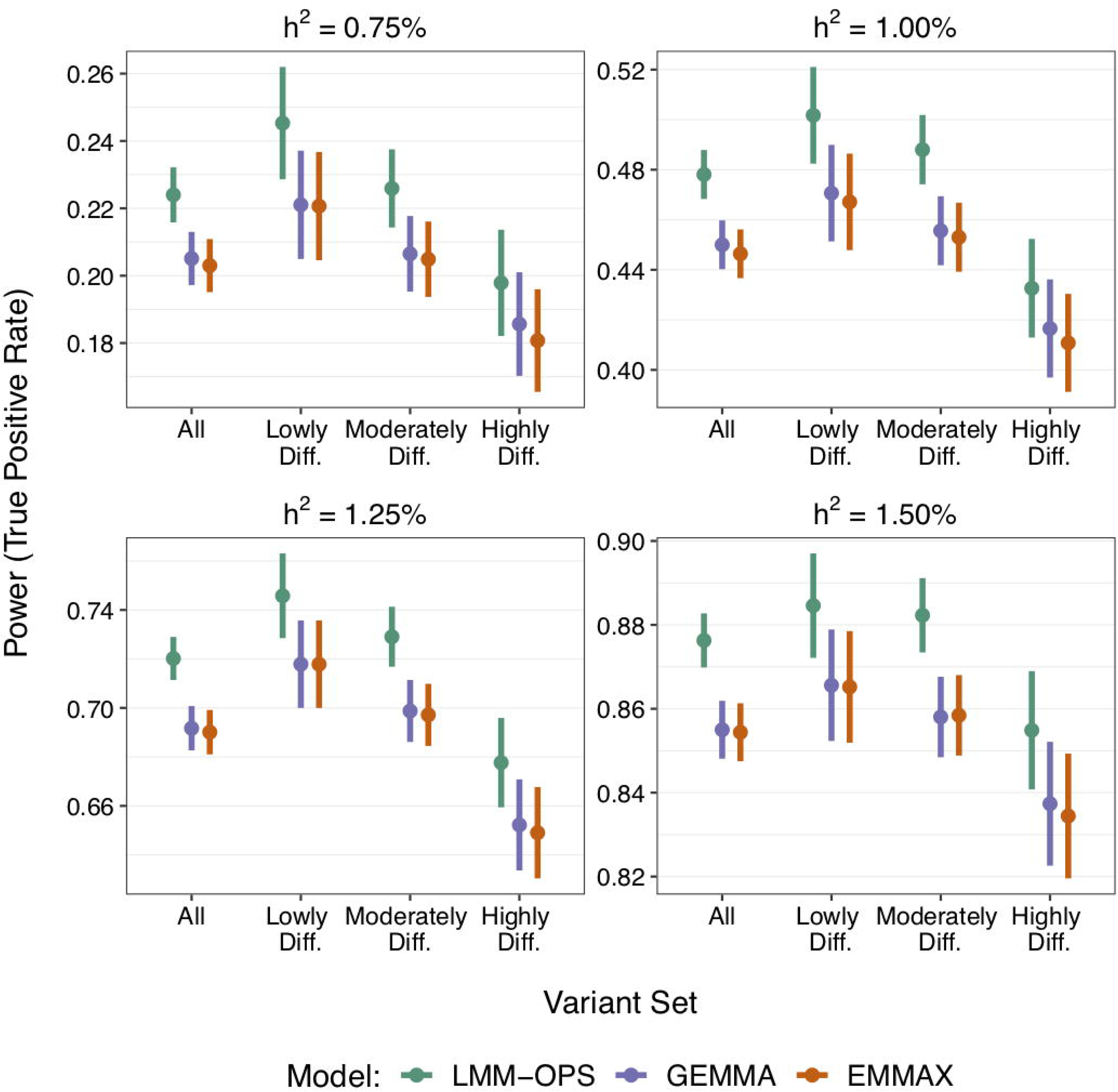
Comparison of power of LMM methods for a phenotype associated with ancestry in the simulation study with admixture and *F*_*ST*_ = 0.15. The power of LMM-OPS, EMMAX, and GEMMA to detect causal SNPs with *h*^*2*^ = 0.75%, 1.00%, 1.25%, and 1.50% is shown across all SNPs as well as within the three classes of SNPs defined by allele frequency differentiation. The points represent the power estimates, and the vertical bars represent the 95% confidence intervals

### GWAS of WBC Count in the WHI SHARe Hispanic Cohort

LMM-OPS, EMMAX, GCTA, and GEMMA all obtained satisfactory genome-wide genomic inflation *λ*_*GC*_ values (Table S3), but the QQ-plots of the –log_10_(*p*-values) from each method appeared to show some early deviation from expectation (Figure 5A). The only SNPs approaching genome-wide significance were on chromosome 1, so we recalculated *λ*_*GC*_ and generated new QQ-plots with chromosome 1 excluded to investigate this deviation. The QQ-plots excluding SNPs from chromosome 1 appeared well behaved for all four methods (Figure 5B), indicating that the early deviation in the genome-wide QQ-plots was due to a large number of associated SNPs on chromosome 1. The corresponding *λ*_*GC*_ values were 1.005, 0.993, 0.993, and 0.994 for LMM-OPS, EMMAX, GCTA, and GEMMA, respectively. We also conducted a GWAS using standard linear regression including the top 6 PCs from PC-AiR as fixed effects. As expected, the test statistics were inflated, giving *λ*_*GC*_ = 1.045 when excluding chromosome 1. This inflation was most likely due to unaccounted for familial relatedness in the sample^20,24^.

**Figure 5.**
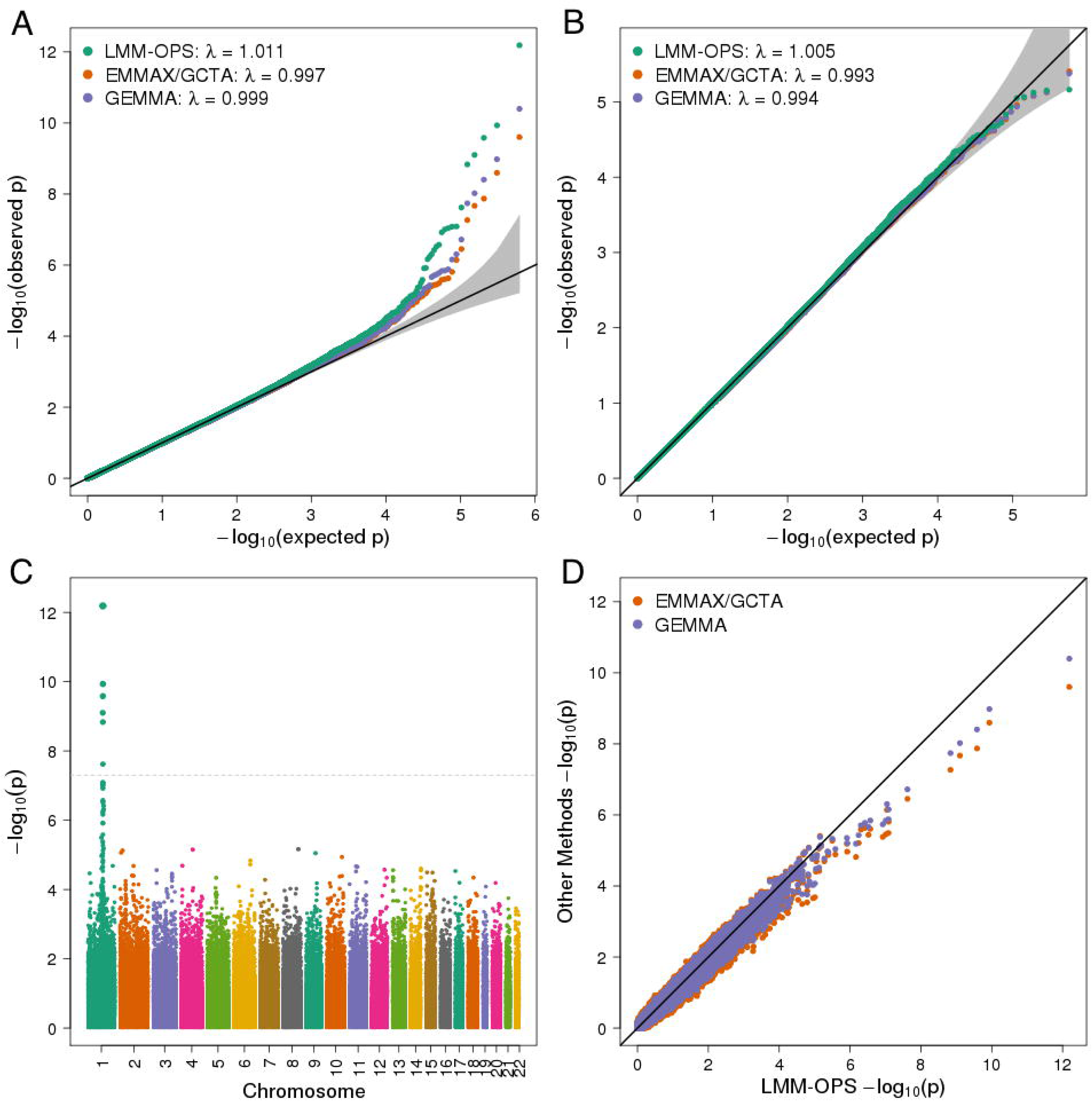
Association testing results for white blood cell (WBC) count in the Hispanic cohort of the WHI-SHARe study. QQ-plots for each of the LMM methods with (A) all autosomal SNPs, and (B) autosomal SNPs excluding chromosome 1. (C) Manhattan plot of the –log_10_(*p*-values) from LMM-OPS. (D) Direct comparison of –log_10_(*p*-values) for all autosomal SNPs from LMM-OPS to each of the other LMM methods. The EMMAX and GCTA results are presented together, as they were nearly identical and could not be distinguished in the figures.

The genotype effect size estimates from all four LMM methods were similar on average, however LMM-OPS consistently provided the smallest standard error estimates, and thus a more efficient association test (Figure S3). The Manhattan plot of the LMM-OPS –log_10_(*p*-values) shows a strong association signal for WBC count in a region on chromosome 1 (Figure 5C). LMM-OPS attained the highest significance in this region, and all genome-wide significant *p*-values with LMM-OPS were one or more orders of magnitude smaller than those from the competing LMM methods (Figure 5D and Table 2). The most significant SNP on chromosome 1 was rs11265198 (LMM-OPS *p* = 6.49 x 10^−13^; EMMAX *p* = 2.49 x 10^−10^; GCTA *p =* 2.77 x 10^-10^; GEMMA *p* = 4.00 x 10^−11^). In addition, there was one SNP in this region, rs6656586, which attained genome-wide significance with LMM-OPS but did not with any of the other methods.

**Table 2.** Genome-wide significant SNPs on chromosome 1 for WBC count in WHI-SHARe Hispanics.

| SNP ID | Position | *p*-value |  |  |  |  |  |  |  |
| --- | --- | --- | --- | --- | --- | --- | --- | --- | --- |
|  |  | LMM-OPS<sup>a</sup> | LMM-OPS<sup>b</sup> | EMMAX | EMMAX<br>+PCs<sup>c</sup> | GCTA | GCTA<br>+PCs<sup>c</sup> | GEMMA | GEMMA<br>+PCs<sup>c</sup> |
| rs11265198 | 159450517 | 6.49 x 10<sup>-13</sup> | 5.75 x 10<sup>-13</sup> | 2.49 x 10<sup>-10</sup> | 2.45 x 10<sup>-10</sup> | 2.77 x 10<sup>-10</sup> | 3.39 x 10<sup>-10</sup> | 4.00 x 10<sup>-11</sup> | 4.14 x 10<sup>-11</sup> |
| rs2808666 | 159591526 | 1.16 x 10<sup>-10</sup> | 1.07 x 10<sup>-10</sup> | 2.53 x 10<sup>-9</sup> | 3.65 x 10<sup>-9</sup> | 2.75 x 10<sup>-9</sup> | 4.44 x 10<sup>-9</sup> | 1.05 x 10<sup>-9</sup> | 1.70 x 10<sup>-9</sup> |
| rs7534472 | 159500861 | 2.62 x 10<sup>-10</sup> | 2.45 x 10<sup>-10</sup> | 1.34 x 10<sup>-8</sup> | 1.36 x 10<sup>-8</sup> | 1.44 x 10<sup>-8</sup> | 1.63 x 10<sup>-8</sup> | 3.94 x 10<sup>-9</sup> | 4.21 x 10<sup>-9</sup> |
| rs857682 | 158670244 | 7.92 x 10<sup>-10</sup> | 6.84 x 10<sup>-10</sup> | 2.14 x 10<sup>-8</sup> | 2.23 x 10<sup>-8</sup> | 2.28 x 10<sup>-8</sup> | 2.73 x 10<sup>-8</sup> | 9.53 x 10<sup>-9</sup> | 1.07 x 10<sup>-8</sup> |
| rs856065 | 159013653 | 1.47 x 10<sup>-9</sup> | 1.44 x 10<sup>-9</sup> | 5.41 x 10<sup>-8</sup> | 6.55 x 10<sup>-8</sup> | 5.73 x 10<sup>-8</sup> | 7.56 x 10<sup>-8</sup> | 1.82 x 10<sup>-8</sup> | 2.45 x 10<sup>-8</sup> |
| s6656586 | 159013653 | 2.40 x 10<sup>-8</sup> | 2.14 x 10<sup>-8</sup> | 3.53 x 10<sup>-7</sup> | 3.92 x 10<sup>-7</sup> | 3.69 x 10<sup>-7</sup> | 4.61 x 10<sup>-7</sup> | 1.91 x 10<sup>-7</sup> | 2.30 x 10<sup>-7</sup> |

The *p*-values are presented for all six SNPs reaching genome-wide significance for WBC count with any of the LMM methods.
<sup>a</sup>LMM-OPS when using the PC-Relate ancestry-adjusted empirical GRM; <sup>b</sup>LMM-OPS when using the centered only ancestry-adjusted empirical GRM. <sup>c</sup>The results labeled “+PCs” are from each of the respective LMM methods when the top 6 PCs from PC-AiR are included as fixed effect covariates

The most significant SNP, rs11265198, is near the Duffy Antigen Receptor for Chemokines (DARC) gene. An African derived regulatory variant in DARC, rs2814778, was previously found to associate with lower WBC count in African Americans^22,23^. Genotype dosages for rs2814778 were imputed using MaCH-Admix^32^ in the WHI-SHARe Hispanics and tested for association using LMM-OPS. The *p*-value for rs2814778 was 1.89 x 10^−18^, providing a stronger signal than any of the directly genotyped SNPs tested for association with WBC count. Additionally, we re-ran a GWAS with LMM-OPS conditional on rs2814778, and all of the previously identified genome-wide significant SNPs on chromosome 1 became non-significant (Figure S4), indicating that these SNPs were tagging this regulatory variant. Thus, we were able replicate and generalize^21^ to Hispanics/Latinos the previously identified association in African Americans for WBC count and this regulatory variant in the DARC gene.

### Importance of Orthogonality

The key feature of LMM-OPS that differentiates it from existing LMM methods for GWAS is the partitioning of sample structure into separate orthogonal components due to recent and distant genetic relatedness. For each of the LMM-OPS analyses presented here, the PC-Relate kinship coefficient matrix, which is calculated conditionally on PCs, was used as the ancestry-adjusted GRM to account for recent genetic relatedness. However, this matrix is actually only approximately orthogonal to the PCs used to account for distant genetic relatedness (see Methods). In order to construct a strictly orthogonal matrix, one could compute an ancestry-adjusted GRM from genotype values that are centered conditional on PCs, without any scaling. Nevertheless, we consistently find that the entries in either of these ancestry-adjusted GRMs are very highly correlated across all pairs of individuals, and the resulting LMM-OPS association test statistics and *p*-values are nearly identical when using either matrix (Figures S5 – S7 and Table 2). Due to this observation, we typically recommend using the PC-Relate kinship coefficient matrix in practice, as it has the advantage of biological interpretability of the matrix elements as kinship coefficient estimates.

To further explore the impact of using an empirical GRM that is orthogonal to the PCs included in the analysis as fixed effect covariates, we re-ran the EMMAX, GCTA, and GEMMA analyses of WBC count, but included the same 6 PCs that were used in the LMM-OPS analysis as fixed effect covariates, which corresponds to an approach that was previously proposed for associating testing in structured samples with unusually differentiated SNPs^15,26^. In these analyses, the PCs and the GRM were both adjusting for the population structure. The results from these models were very similar to those from the corresponding models without PCs (Figure S8 and Table 2). These results demonstrate that the efficiency gain achieved with LMM-OPS cannot be replicated by simply including ancestry representative PCs in an LMM with a standard empirical GRM. Partitioning sample structure due to population stratification and recent genetic relatedness into separate, orthogonal components, as is done with LMM-OPS, is essential for improved efficiency in association testing.

### Computation Time

We compared the computation time required by LMM-OPS, EMMAX, GEMMA (v0.94), and GCTA (v1.24.7) to perform the association analysis of WBC count at all 616,556 autosomal SNPs for the 3,551 women in the WHI-SHARe sample. The computation times required to construct the relationship matrices were not included in this assessment since all of the LMM methods can use a pre-computed GRM, and each GRM has similar computational complexity^10^. Each method was run using a single core 2.4 GHz Intel Xeon E5-2630L processor with 128 GB of RAM. LMM-OPS is implemented in R, and was run using R v3.2.0 configured to use the BLAS and LAPACK libraries within the Intel Math Kernel Library (MKL). LMM-OPS was substantially faster than the other methods, which took at least twice as long, and up to over seven times as long to perform the analysis. The computation time for LMM-OPS to complete the analysis was 20.4 minutes (0.34 hours). In comparison, the computation time for GCTA was 44.6 minutes (0.74 hours), for GEMMA was 1.84 hours, and for EMMAX was 2.43 hours.

## Discussion

Genetic association studies involving ancestrally diverse populations from around the world have recently become more common as there is increased interest in both identifying novel, population specific variants that underlie phenotypic diversity and generalizing associations across populations. Confounding due to ancestry is a serious concern for genetic association studies since different ethnic groups often share distinct dietary habits and other lifestyle characteristics that lead to many traits of interest being correlated with ancestry. Linear mixed models (LMMs) have become the go-to approach for genome-wide association testing of quantitative traits, as they are both computationally fast and statistically powerful^10^. Furthermore, it has been reported that LMMs are effective at controlling type I error rates in samples with relatedness and population structure, as they tend to provide acceptable genomic control inflation factors. However, we demonstrated through simulation studies that existing implementations of LMMs for GWAS can provide systematically biased test statistics in samples with population stratification when phenotypes are correlated with ancestry. Interestingly, we found that the miscalibration of test statistics from widely used LMM approaches is a problem that can occur when samples descend from either highly divergent inter-continental populations or closely related intra-continental populations. Additionally, the problem manifests in the presence of both discrete population substructure and ancestry admixture.

Incorrect calibration of association test statistics from existing LMM methods arises because they use an empirical GRM calculated as a genome-wide average genotype covariance to account for the entire sample genealogy, including both distant population structure and recent familial relatedness. However, as we demonstrated using genotype data from release 3 of phase III of HapMap, allele frequency differences among human populations vary greatly by SNP across the genome. As a consequence, the strength of the genetic covariance due to ancestry also varies by SNP across the genome, and existing LMM methods can provide inadequate correction for population stratification. The ancestry correction provided by these methods is of appropriate size only for SNPs with typical allele frequency differences between populations, while SNPs with relatively larger or smaller allele frequency differences receive an under- or over-correction, respectively. This result is contrary to what has previously been suggested^10^, that SNPs with larger allele frequency differences between populations receive a larger correction from these LMM methods. Notably, the deflation of test statistics for SNPs with similar allele frequencies across populations, and the inflation of test statistics for SNPs that are highly differentiated across populations, balances out across the genome, typically leading to a genomic control inflation factor near 1 and QQ-plots that look acceptable when considering all SNPs genome-wide, which likely contributed to why this phenomenon had not been previously reported.

To address this issue, we developed LMM-OPS, which orthogonally partitions the genealogical structure among sampled individuals into two separate components. Ancestry-representative vectors are included as fixed effects in the mean model of LMM-OPS to account for population stratification, and an ancestry-adjusted empirical GRM that is orthogonal to the ancestry-representative vectors is used to model recent familial relatedness as a random effect. In simulation studies with real and simulated genotype data, we demonstrated that the LMM-OPS testing procedure provides well-calibrated association test statistics at *all* SNPs genome-wide, regardless of the distribution of allele frequency differences among the underlying populations. In addition to providing better protection against false positives, we also demonstrated that, compared to existing LMM methods, LMM-OPS provides a more efficient test with improved power to detect true SNP-phenotype associations. This increase in power holds even at highly differentiated SNPs, for which existing LMM methods provide systematically inflated test statistics resulting in an inflated type-I error rate.

We also compared the performance of LMM-OPS to existing implementations of LMMs through a GWAS analysis of white blood cell (WBC) count in the Hispanic cohort of the WHI SHARe study. This cohort contains multi-way continental ancestry admixture as well as cryptic familial relatedness. All four methods gave similar genotype effect size estimates at SNPs for WBC count, but LMM-OPS was the most efficient, providing consistently smaller standard errors and genome-wide significant *p*-values that were more significant, by one or more orders of magnitude, than EMMAX, GEMMA, and GCTA. Using LMM-OPS, we were able to replicate generalize to this Hispanic American population (*p* = 1.89 x 10^−18^) the association at regulatory variant rs2814778 in the Duffy Antigen Receptor for Chemokines (DARC) gene previously identified in African Americans. Because of natural selection, rs2814778 is highly differentiated between African and European ancestral populations, likely due to a protective effect against *P. vivax* malaria^34,35^. Furthermore, through a conditional analysis including rs2814778 as a covariate, we were able to demonstrate that other genome-wide significant associations in this region on chromosome 1 could be explained by LD with this particular variant.

In the implementation of LMM-OPS presented here, we utilized ancestry-representative principal components (PCs) from PC-AiR to adjust for population structure and construct an orthogonal ancestry-adjusted empirical GRM to account for relatedness. Alternatively, vectors of estimated individual admixture proportions from model-based methods such as ADMIXTURE^36^ or FRAPPE^37^ could be used in place of PCs. We also performed the LMM-OPS analysis of WBC count in the Hispanic American cohort of the WHI-SHARe using model-based estimates of individual ancestry from a supervised ADMIXTURE analysis that included reference population samples from the International Haplotype Map Project (HapMap) and the Human Genome Diversity Project (HGDP)^38^ for European, Native American, African, and East Asian ancestry. The results were nearly identical to the analysis that used PCs (Figure S9). In general, using model-based estimates of ancestry with LMM-OPS is expected to work well, provided that prior assumptions regarding the underlying populations contributing ancestry to the sample are accurate and suitable reference population panels representative of these populations are available for reliable estimation of ancestry. For many genetic studies, however, the underlying ancestral populations may not be completely known or well defined, and misspecification of the ancestral populations will result in less efficient association tests due to biased estimates of individual ancestry from model-based methods. For this reason, we recommend using PCs with LMM-OPS, as they do not rely on strong modeling assumptions.

Numerous LMM approaches for GWAS have been proposed in the past few years, each with small variations on the testing procedure. One variation among these methods lies in SNP selection for constructing the empirical GRM. In each of the analyses performed here, the GRM was constructed from all SNPs genome-wide with sample minor allele frequency (MAF) greater than 1%. Others have suggested that including the SNP being tested, or SNPs in LD with the one being tested, leads to proximal contamination and a loss of power^6,10^. This has led to the development of automated methods for selecting subsets of SNPs to be used in the GRM^6^, as well as the leave-one-chromosome-out (LOCO) approach^10^, where the GRM is constructed from all autosomal SNPs not on the same chromosome as the SNP being tested. While not presented here, LMM-OPS can easily incorporate using a subset of SNPs or a LOCO approach for constructing the ancestry-adjusted GRM. However, it remains unclear as to what the optimal set of SNPs is for obtaining the GRM. Using a subset of SNPs, as is done with FaST-LMM-Select^6^, may improve power, but it may also provide inadequate control of type-I error rates^39^. When there is family relatedness among samples, this remains true even if PCs are included in the model^39^. Similarly, using a LOCO approach, such as that implemented in GCTA-LOCO^10^, may also improve power, but may inadequately account for the effects of SNPs on the same chromosome as the SNP being tested. A hybrid method that uses two GRMs, one constructed from all SNPs, and the other from a selected subset of SNPs, has also been proposed^39^. However, this approach remains susceptible to the systematic inflation/deflation issues illustrated in this work for phenotypes correlated with ancestry. We have demonstrated that LMMs for GWAS should certainly include ancestry-representative vectors as fixed effects to account for population stratification, but exactly which SNPs should be used to construct the empirical GRM in order to optimize power, while still adequately protecting against false positives due to family structure, remains an open area of research.

### Appendix A

#### Sample Structure Inference

Population structure inference for LMM-OPS and for linear regression adjusted with PCs was performed using PC-AiR^25^. PC-AiR used SNPs LD-pruned with an *r*^*2*^ = 0.1 threshold in a sliding 10Mb window for the HapMap and WHI-SHARe analyses, and it used all SNPs in Set 1 for the simulations with admixture. Inference on recent genetic relatedness due to family structure for LMM-OPS was performed using PC-Relate^23^; these kinship coefficient estimates were used to construct the ancestry-adjusted empirical GRM. EMMAX, GEMMA, and GCTA each used the default empirical GRM created by the respective software to infer all sample structure. The GRM for each method was constructed from all of the autosomal SNPs with minor allele frequency (MAF) > 1% in the HapMap and WHI-SHARe analyses, and from all SNPs in Set 1 in the simulations with admixture.

### Appendix B

#### Simulated Genotype Data with Admixture

For each of the three simulated data sets with admixture, allele frequencies for the two populations at all 200,000 SNPs were generated using the Balding-Nichols model^30^. More precisely, for each SNP *s*, the allele frequency *p*_*s*_ in the ancestral population was drawn from a uniform distribution on [0.1, 0.9]. The allele frequency in each population was then drawn from a beta distribution with parameters *p*_*s*_*(1–F*_*ST*_*)/F*_*ST*_ and *(1–p*_*s*_*)(1–F*_*ST*_*)/F*_*ST*_, where the quantity *F*_*ST*_ is Wright’s measure of genetic distance between populations^27^. The vector **p**_*s*_ contains the allele frequencies at SNP *s* for each population. An individual’s vector of ancestry proportions from each of the two populations can be represented by **a**_*i*_ ^*T*^ = (*a*,1– *a*). The parameter *a* was drawn from a beta distribution with mean 0.3 and s.d. 0.1 for one third of unrelated individuals and pedigree founders, a beta distribution with mean 0.7 and s.d. 0.1 for another third, and a uniform distribution on [0,1] for the remaining third. Within any given pedigree, every founder had *a* drawn from the same distribution. As a result, individuals in a pedigree for which founder ancestry proportions were drawn from either of the beta distributions had similar ancestry to each other, which could be viewed as a type of ancestry-related assortative mating. Ancestry proportions for pedigree descendants were calculated as the average of their parents’ ancestry proportions. Genotypes for unrelated individuals and pedigree founders were randomly drawn from a 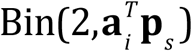 distribution, and alleles were passed down the pedigree to generate genotypes for pedigree descendants (including the cousin pairs).

#### Simulated Phenotypes

For the simulations used to assess behavior at null SNPs, each replicate of the heritable phenotype whose mean depended on genome-wide ancestry (or population membership) was generated according to the model

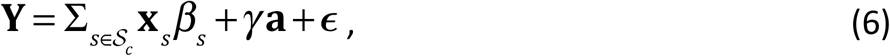

where 𝒮_*c*_ is a set of 100 randomly selected causal SNPs, **x**_*s*_ is the vector of genotype values for all individuals at SNP *s* ∈ 𝒮_*c*_ with effect size *β*_*s*_ chosen using sample allele frequencies so that SNP *s* explains 0.1% of the phenotypic variability, **a** is the vector of ancestry proportions for all individuals with effect size *γ,* and **ε**_*i*_ ∼ N(0,1) is independent random noise. In the simulations using the HapMap genotype data, SNPs for 𝒮_*c*_ were selected randomly from chromosome 1, **a** was 1 for an individual in the CEU population and 0 otherwise, and *γ* = 1. In the simulations using the simulated genotype data with admixture, SNPs for 𝒮 _*c*_ were randomly chosen from Set 2, **a** was an individual’s ancestry proportion from population 1, and *γ* = *2.*

For the simulations used to evaluate detection of causal SNPs, each of the causal SNP-phenotype pairs was generated according to the model

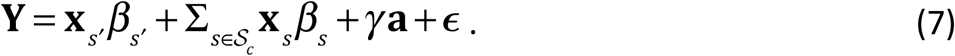

This is the same model as in Equation (6) but with one additional causal SNP of interest, *s’*. In the simulations using the simulated genotype data with admixture, *s’* was selected at random from Set 2. The effect size *β*_*s*’_ for SNP *s’* was chosen to explain a pre-specified proportion of the phenotypic variability, denoted 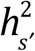 and 10,000 replicate SNP-phenotype pairs were simulated for each choice of 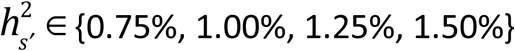.

## Supplemental Data

Supplemental data include 9 Figures and 3 Tables.

## Acknowledgements

This work was supported in part National Institutes of Health (NIH) grants P01 GM 099568 (to T.A.T.), K01 CA148958 (to T.A.T), R01 HG001645 (to M.S.M.), and R01 HL116446 (to A.P.R. and T.A.T.). The WHI program is funded by the National Heart, Lung, and Blood Institute, National Institutes of Health, U.S. Department of Health and Human Services through contracts HHSN268201100046C, HHSN268201100001C, HHSN268201100002C, HHSN268201100003C, HHSN268201100004C, and HHSN271201100004C. Funding for WHI SNP Health Association Resource (WHI-SHARe) genotyping was provided by NHLBI contract N02-HL-64278.

## Web Resources

LMM-OPS is implemented in the R language and is freely available from Bioconductor as part of the GENESIS package (http://www.bioconductor.org/packages/release/bioc/html/GENESIS.html).

